# Phosphorylation of a *Toxoplasma gondii* tyrosine transporter by calcium-dependent kinase 3 is important for parasite fitness

**DOI:** 10.1101/339671

**Authors:** Bethan A. Wallbank, Caia S. Dominicus, Malgorzata Broncel, Nathalie Legrave, James I. MacRae, Henry M. Staines, Moritz Treeck

## Abstract

*Toxoplasma gondii* parasites rapidly exit their host cell when exposed to calcium ionophores. The calcium-dependent protein kinase 3 (*Tg*CDPK3) was previously identified as a key mediator in this process, as *Tg*CDPK3 knockout (Δ*cdpk3*) parasites fail to egress in a timely manner. Phosphoproteomic analysis comparing WT with Δ*cdpk3* parasites revealed changes in the *Tg*CDPK3-dependent phosphoproteome that included proteins important for regulating motility, but also metabolic enzymes, indicating that *Tg*CDPK3 controls processes beyond egress. Here we have investigated a predicted direct target of *Tg*CDPK3, a putative transporter of the major facilitator superfamily (MFS) and show that it is rapidly phosphorylated after induction of calcium signalling. Conditional knockout (KO) of the transporter reveals an essential role in the lytic cycle during intracellular growth with a transcriptome signature of amino acid-starved parasites. Using a combination of metabolomics and heterologous expression, we confirmed a primary role in tyrosine import. Complementation with phosphorylation site mutants shows that phosphorylation of serine 56 (S56) by *Tg*CDPK3 gives the parasites a growth benefit in competition assays. Collectively, these findings validate an important, albeit non-essential role for *Tg*CDPK3 in the regulation of metabolic processes, in addition to motility.

**Author summary:** *Toxoplasma gondii* is an obligate intracellular parasite. To survive and spread throughout the host it must repeatedly infect, replicate within and exit, host cells. These recurring cycles of infection and egress rely on signalling pathways that allow the parasites to sense and respond rapidly to their environment. While some key kinases and secondary messengers within these pathways have been identified, functional analysis of non-kinases has been very limited. This is especially true for candidates that are not predicted to play a role in active motility or are not known to function in established signalling pathways. Here we have followed up on an unexpected target of the *T. gondii* calcium-dependent kinase 3 (*Tg*CDPK3), a plant-like calcium dependent kinase, that was previously shown to play an important role in calcium-mediated exit from the host cell. We show that, in addition to controlling motility of the parasite (as previously shown), *Tg*CDPK3 phosphorylates an essential tyrosine transporter in the plasma membrane. Mutational analysis of the phosphorylation sites demonstrates an important role in maintaining parasite fitness, thus demonstrating that *Tg*CDPK3 plays a pleiotropic role in controlling both egress and metabolism.

## Introduction

The fast growing tachyzoite stage of the protozoan parasite *Toxoplasma gondii* requires cycles of host cell invasion, replication, and lysis for its successful proliferation within the host. Each step of this lytic cycle involves tightly regulated signalling pathways, the intricacies of which remain largely unknown. Paramount to parasite survival is the ability to sense and respond to changes in the environment for which the divalent calcium ion (Ca^2+^) acts as an important secondary messenger (1). Changes in free intracellular [Ca^2+^]_i_ levels, via release of Ca^2+^ from organellar Ca^2+^ stores, can be induced by the addition of Ca^2+^ ionophores, such as A23187 or phosphodiesterase inhibitors (2,3). Ca^2+^ flux regulates key processes including secretion of micronemes prior to host cell entry (4), parasite motility (5), and host cell egress (6) and invasion (7). Inversely, these processes can all be inhibited by Ca^2+^ immobilisers or chelators, such as BAPTA-AM (5,8–10). Ca^2+^ release leads to the activation of Ca^2+^ binding proteins such as calmodulins, calcineurin B-like kinases and calcium-dependent protein kinases (CDPKs). *T. gondii* calcium-dependent protein kinase 3 (*Tg*CDPK3), for example, has been implicated in the regulation of ionophore induced egress, IIE (i.e. the rapid exit of tachyzoites from a host cell on addition of ionophore) and ionophore induced death, IID (i.e. the loss of infectivity of EC parasites after prolonged exposure to ionophore) (9). *TgCDPK3* KO (Δ*cdpk3*) (11), mutants (9), and chemically inhibited *Tg*CDPK3 lines (12) all show a deficiency in IIE and IID. *Tg*CDPK3 is a serine/threonine kinase belonging to a large family of CDPKs also found in plants and ciliates, but absent in humans. It is anchored to the parasite plasma membrane, via N-terminal myristoylation and palmitoylation motifs (11–13), facing the lumen of the parasite. Like all CDPKs, *Tg*CDPK3 possesses a C-terminal calmodulin-like domain consisting of 4 EF hands, known as the CDPK activation domain, as well as upstream autoinhibitory and catalytic domains (14). Binding of Ca^2+^ to the EF hands causes a structural rearrangement that frees up the active site of the kinase domain, allowing for substrate phosphorylation (15,16). A quantitative phosphoproteome study revealed 156 phosphorylation sites that are differentially phosphorylated between WT and *Tg*CDPK3 mutant parasites (17). The *Tg*CDPK3-dependent phosphoproteome includes phosphorylation sites on proteins involved in parasite motility, such as the cyclase-associated protein and myosin A (Myo A), but also, and perhaps surprisingly, those involved in metabolic processes such as the α-ketoacid dehydrogenase (BCKDH) subunit, E1α, required for the breakdown of branched-chain amino acids (BCAAs) and conversion of pyruvate to the TCA driver acetyl-CoA (18). The link to proteins not obviously involved in egress and motility, as well as changes in the phosphoproteome regardless of the presence of ionophore, suggests that *Tg*CDPK3 function extends beyond egress.

The phosphorylation site that appeared to have one of the most marked reductions in phosphorylation state in *Tg*CDPK3 mutants compared to WT parasites (17), is situated within a putative transporter of the MFS family (TGGT1_257530, named ApiAT5-3 as per (19)) that has moderate homology to a BCAA transporter. Given the additional evidence from the phosphoproteomic dataset that *Tg*CDPK3 putatively regulates BCAA catabolism via BCKDH, we hypothesised that *Tg*CDPK3 might be involved in BCAA uptake and metabolism via phosphorylation of ApiAT5-3 in addition to regulating motility. Here, we have assessed the function of ApiAT5-3 and the role of *Tg*CDPK3-mediated phosphorylation. We show that ApiAT5-3 is rapidly phosphorylated at serine 56 (S56) during the first minute of induced egress. Using a conditional KO approach, we show that ApiAT5-3 is essential, and that deletion leads to a delayed death phenotype that is accompanied by a transcriptional response relating to translational stress. In growth competition assays performed with parasite lines that rely on phosphomutants or phosphomimetics, we show that phosphorylation of S56 appears to be important, but not essential, for parasite fitness. Finally, using a combination of metabolic analysis and heterologous expression in *Xenopus laevis* oocytes we confirm that ApiAT5-3 transports tyrosine but has only limited capacity to transport BCAAs. This data confirms that *Tg*CDPK3 phosphorylates several targets in its vicinity, controlling diverse processes at the plasma membrane and thus contributing to a range of biological processes in the parasite.

## Results

### ApiAT5-3 is located at the parasite periphery and phosphorylated during ionophore induced egress in a *Tg*CDPK3-dependent manner

ApiAT5-3 was previously identified to be phosphorylated at Serine 56 in a *Tg*CDPK3-dependent manner (17). BLAST analysis using the Transporter Classification Database (http://www.tcdb.org/) predicts that ApiAT5-3 possesses a modest level of homology to a BCAA transporter. This was interesting, as deletion of *TgCDPK3* was previously shown to lead to upregulation of the BCKDH complex (17), involved in BCAA catabolism. This indicated that *Tg*CDPK3 may directly control BCAA transport by phosphorylating ApiAT5-3.

Topology prediction (https://embnet.vital-it.ch/software/TMPRED_form.html (20)) places the N-terminal regions of ApiAT5-3 at the luminal side of the parasite, potentially allowing for direct interaction with *Tg*CDPK3, which also localises to the plasma membrane (Fig. 1A). ApiAT5-3 contains several phosphorylation sites at its N-terminus, of which S56 was the only one previously identified as being *Tg*CDPK3-dependent (Fig. 1B, upper panel). It is entirely plausible, however, that kinases other than *Tg*CDPK3 act during egress to phosphorylate additional residues on the ApiAT5-3 N-terminus. To investigate this, we queried a dataset recently generated in our laboratory in which we have quantified, using tandem-mass-tag technology (21), phosphorylation site abundance on *T. gondii* proteins across 4 time points (0, 15, 30 and 60 s) following ionophore-treatment (Caia Dominicus, in preparation). From the ~850 phosphorylation sites that are phosphorylated or de-phosphorylated during egress, we identified S56 of ApiAT5-3 as increasingly phosphorylated over time (Fig. 1B, lower panel). We also identified several proteins already known to be more phosphorylated in response to Ca^2+^ signalling including MyoA, Myosin F and DrpB (17,22,23). None of the other phosphorylation sites on the ApiAT5-3 N-terminus increased in phosphorylation state prior to, or during egress. However, S14 of ApiAT5-3 was dephosphorylated during ionophore-treatment. Collectively these data indicate that S56 is phosphorylated in a *Tg*CDPK3-dependent manner upon Ca^2+^ stimulation, and that a phosphatase is acting on S14 during the same period, while the other phosphorylation sites appear unaffected.

**Fig 1.**
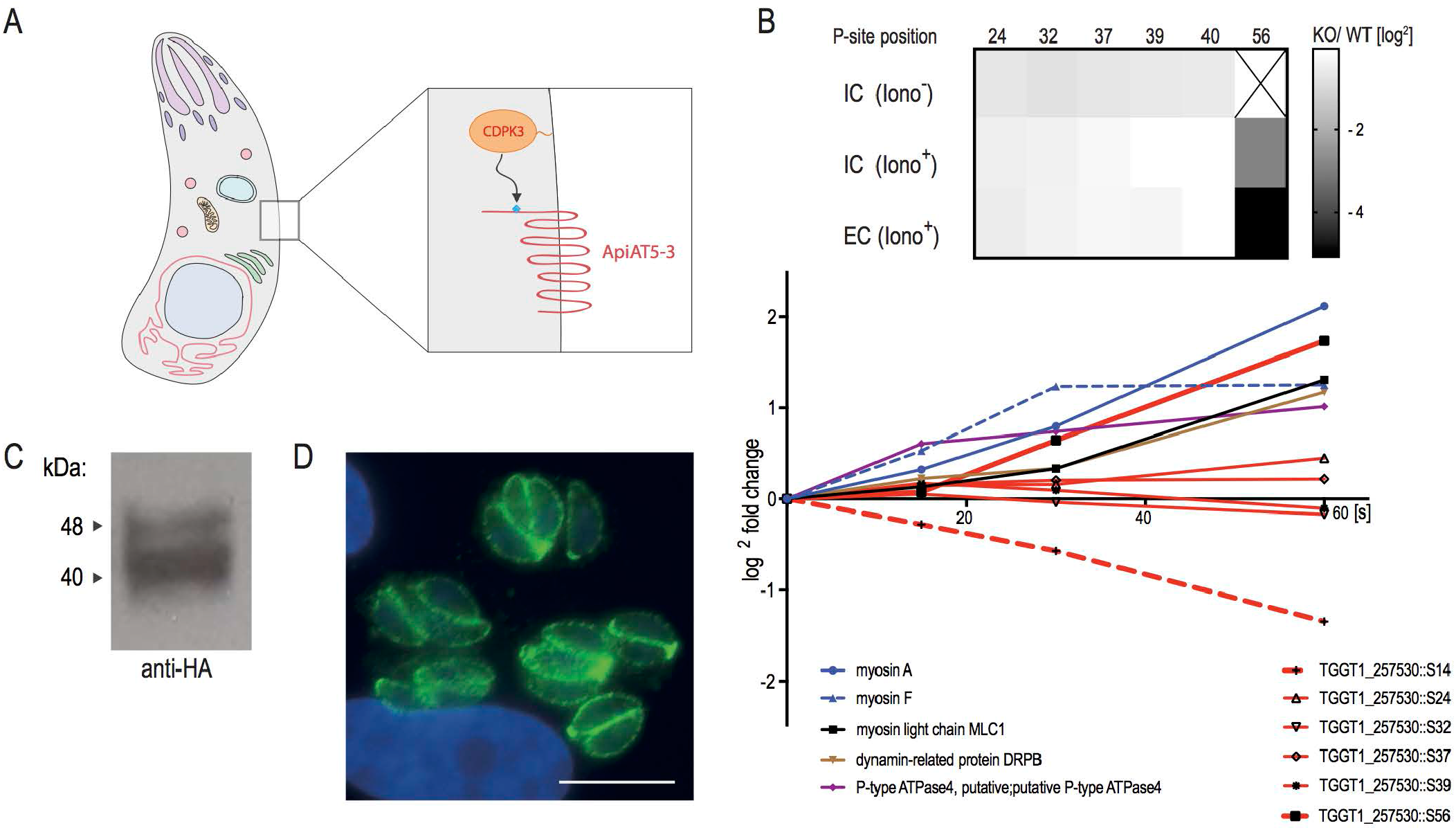
ApiATP5-3 localises to the plasma membrane and is phosphorylated at serine 56 upon ionophore treatment. **(A)** Topology prediction of ApiAT5-3. Serine 56 is predicted to be located on the luminal side of the parasite **(B)** Quantification of the phosphorylation state of residues in the ApiAT5-3 N-terminus in *TgCDPK3* KOs and during ionophore-induced egress (from (17)). Upper panel: The heatmap shows differential phosphorylation of S56 in *Tg*CDPK3 mutants compared to WT parasites, but not any other of the identified phosphorylation sites. Intracellular (IC) and extracellular (EC) parasites with and without 1 μM ionophore (iono). P-site = phosphorylation site. Numbers represent residue position. Black ‘x’ = phosphorylation site not identified. Fold changes are log^2^. Bottom panel: Change in relative phosphorylation of ApiAT5-3 and proteins with previously described ionophore-dependent phosphorylation sites, measured after addition of 8 μM ionophore over 60 s. Numbers after the identifier represent the phosphorylation site quantified. **(C)** ApiAT5-3 was detected by Western blot analysis of ApiAT5-3::HA cell lysate using anti-HA antibody. **(D)** IFA of ApiAT5-3::HA expressing parasites shows that ApiAT5-3 localises to the periphery of the intracellular tachyzoite. Green = HA. Scale bar 10 μm.

To localise ApiAT5-3 in *T. gondii* parasites we generated an RH parasite line (ApiAT5-3::HA) that expressed a C-terminally HA-epitope tagged version of ApiAT5-3 under control of its endogenous promotor (i.e. 1000 bp upstream of the start ATG). Western Blotting confirmed expression of a 42 KDa protein close to the predicted size (56 KDa) (Fig. 1C) and immunofluorescence assays (IFA) showed ApiAT5-3 predominantly at the periphery of the parasite (Fig. 1D). No ApiAT5-3::HA could be detected in nascent daughter cells, a hallmark of most inner membrane complex proteins. Together, these data suggest that ApiAT5-3 localises, like *Tg*CDPK3, to the plasma membrane, and thus, could be a *bona fide* target of *Tg*CDPK3 *in vivo*.

### *apiAT5-3* deletion causes delayed parasite death

ApiAT5-3 depletion is predicted to have a high fitness cost (Toxo DB 7.1 (24)). Accordingly, we generated a conditional KO using the dimerisable cre recombinase (DiCre) strategy. We replaced the endogenous copy of *apiAT5-3* with a recodonised version in RH Δ*ku80*^DiCre^ parasites, by double homologous recombination, using a double-guide strategy (25) (Fig. 2A). We initially placed a loxP site adjacent to the Kozac sequence of ApiAT5-3 but were unable to obtain correct integration. We hypothesised that the loxP sequence might be interfering with promotor elements and moved it 100 and 200 bp upstream of the predicted start ATG, respectively. Both of these constructs correctly integrated into the genome. Subsequent analyses were performed with the resulting ApiAT5-3_loxP, with the loxP at ATG −100bp. Integration was confirmed by PCR amplification (Fig. 2B, left panel). To test whether ApiAT5-3 is an essential gene we treated parasites for 4h with either rapamycin (RAP) or DMSO. PCR analysis showed a near complete excision of the floxed gene (Fig. 2B, right panel). Correct excision of the ApiAT5-3 open reading frame resulted in YFP positive parasites that could be readily distinguished from WT by microscopy (Fig. 2C). Upon performing plaque assays it became evident that RAP, but not DMSO-treatment, resulted in a complete block in plaque formation (Fig. 2D). A small number (<0.5%) of plaques could be identified in RAP-treated cultures, however, parasites contained in these plaques where YFP(−), indicating that they arose from non-excised parasites (data not shown). Over time these non-excised parasites within the RAP-treated population outgrew the KOs (Fig. S1), further reinforcing the fact that ApiAT5-3 is essential for parasite survival. These non-excising parasites (termed ApiAT5-3_loxP^dDiCre^), which presumably possess a non-functioning diCre recombinase, were isolated and used as controls for subsequent experiments, as detailed below.

**Fig 2.**
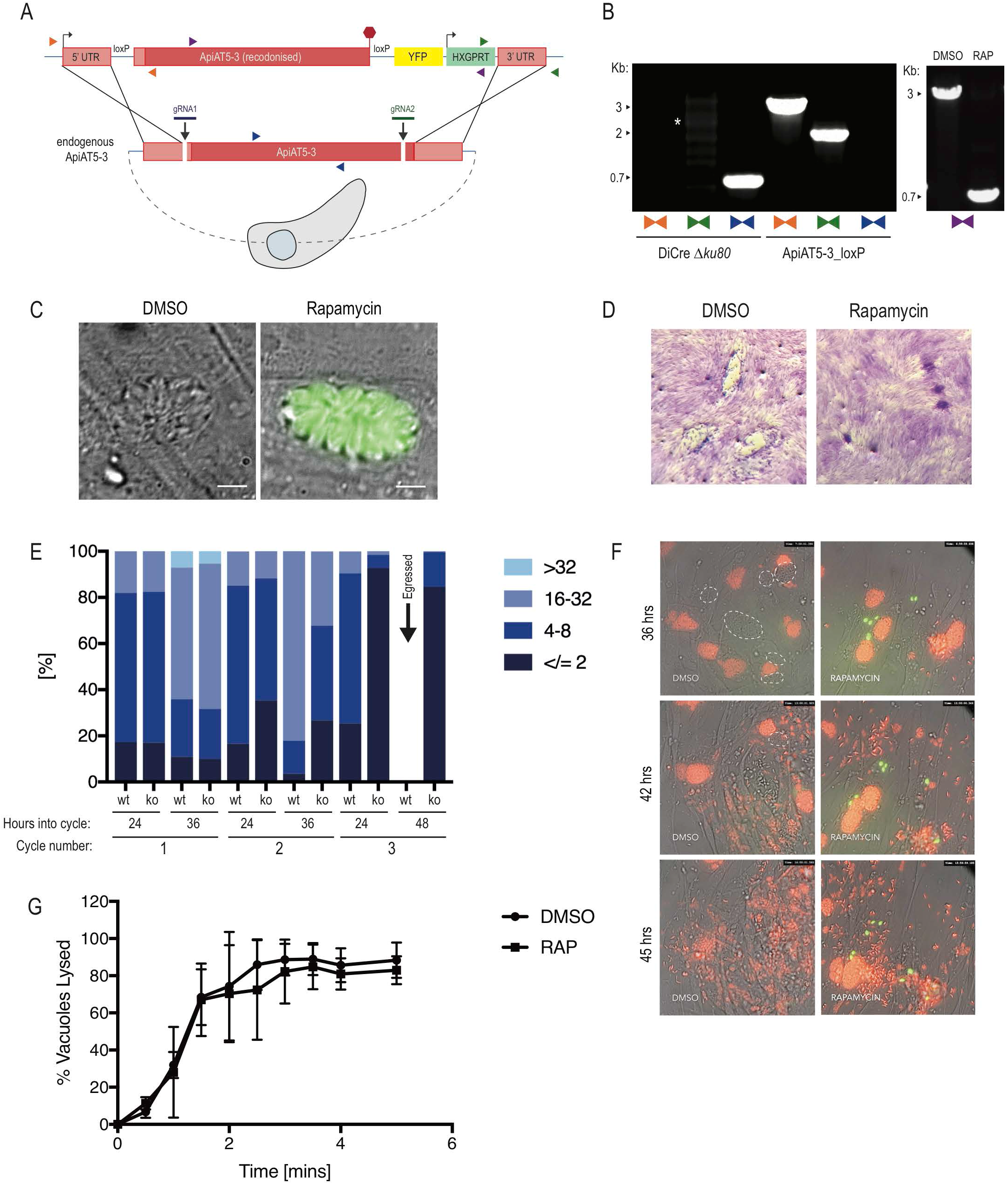
ApiAT5-3 is essential for parasite proliferation. **(A)** Generation of the ApiAT5-3_loxP line using CRIPSPR/Cas9 to increase site-directed integration. Protospacer adjacent motif (PAM) indicated by black arrows. Primer pairs represented by coloured triangles. **(B)** Left panel: PCR analysis shows correct integration of the ApiAT5-3_loxP construct at both the 3’ and 5’ ends and a loss of WT *apiAT5-3* at the endogenous locus. White * = non-specific bands. Right panel: Addition of RAP leads to correct recombination of the loxP sites. **(C)** Fluorescent microscopy of ApiAT5-3_loxP parasites 24 hrs after addition of DMSO or RAP. Scale bar 5 μm **(D)** Plaque assay showing loss of plaquing capacity of ApiAT5-3_loxP parasites upon RAP-treatment. **(E)** Parasite per vacuole number shown as mean %. n=3 **(F)** Stills from live video microscopy at 36, 42 and 45 hrs into 3^rd^ lytic cycle post RAP-treatment. Red = RH Tom, dashed white line = WT ApiAT5-3_loxP, green = *apiAT5-3* KO. **(G)** Ionophore induced egress assay showing no significant difference between DMSO and RAP-treated ApiAT5-3_loxP (at 30 hrs into lytic cycle 2 post DMSO/RAP-treatment). Statistical analysis using multiple comparison 2-way ANOVA, n = 3.

To visualise at which time points ApiAT5-3 is important, we followed replication of live RAP-treated ApiAT5-3_loxP parasites over 3 lytic cycles, where each lytic cycle is defined as growth over 36 hrs, before passage into a fresh culture dish containing host cells. This analysis revealed that in the first cycle, RAP-treated (*apiAT5-3* KO) parasite numbers and replication rate remained comparable to DMSO-treated (WT) parasites (Fig. 2E). However, by 36 hrs into the second replicative cycle there was a 60.7% decrease in the number of vacuoles with more than 16 parasites compared to the DMSO control. By the end of the 3^rd^ cycle the *apiAT5-3* KO parasites consisted largely of 2 or fewer parasites/vacuole, even after the WT had successfully egressed (48 hrs into cycle). To better identify phenotypic consequences of *apiAT5-3* deletion, we followed replication over time using live-video microscopy. We started recording 29 hrs into the third lytic cycle post RAP-treatment, by which time *apiAT5-3* KO parasites display a marked growth defect. To facilitate a more accurate comparison between *apiAT5-3* KO and WT parasites, tdTomato expressing RH parasites (RH Tom) were spiked into the imaging wells at a 1:1 ratio. These analyses revealed that *apiAT5-3* KO does not lead to early egress or an inability to invade, but rather a lack of replication, often with the ability to undergo a first division, but failing to go beyond 2 parasites/vacuole (Fig. 2F, Movies S1A and B).

As we showed that ApiAT5-3 is phosphorylated directly after ionophore-treatment (Fig. 1B), we postulated that it may be required for ionophore induced egress. To assess this, we performed egress assays of the DMSO- and RAP-treated lines in the presence of 8 μM Ca^2+^ ionophore. However, there was no significant difference between the KO and WT (Fig. 2G) suggesting that phosphorylation of ApiAT5-3, in response to elevated Ca^2+^ levels, plays a role in processes other than egress.

### *apiAT5-3* KO parasites display a transcriptional response related to amino acid starvation

Deletion of a transporter may lead to up-regulation of alternative transporters or may manifest as a stress response that carries a detectable signature. To investigate this, we measured transcript levels using RNAseq, comparing RAP-treated ApiAT5-3_loxP with ApiAT5-3_loxP^dDiCre^ parasites, which, as mentioned previously, do not excise the endogenous locus, even when treated with RAP. RNA was isolated in biological triplicate at 4 hrs post RAP-treatment, the time point at which we did not expect to see major changes in the transcriptome, and 60 hrs post treatment, by which time point the RAP-treated ApiAT5-3_loxP parasites are still viable but start to display a growth defect. Indeed, at 4 hrs transcripts from the *apiAT5-3* gene were only slightly reduced in the RAP-treated ApiAT5-3_loxP parasites compared to the RAP-treated ApiAT5-3_loxP^dDiCre^ parasites (17.1%). In contrast, at 60 hrs post RAP-treatment, a 64.7% reduction was observed (Fig. 3A). Unexpectedly, only 435 transcripts showed a statistically significant differential expression between the WT and *apiAT5-3* KO parasites at the 60 hrs time point, compared to the 4 hrs time point, indicating a modest transcriptional response to *apiAT5-3* deletion (Fig. 3B). GO-term analysis of the differentially transcribed genes showed most enrichment (5.41-fold) for genes important for translation. Among this enriched group, these genes encode almost exclusively genes for ribosomal proteins (Fig. 3C, Table S3). No single transporter was specifically up-regulated, indicating that there is no rapid transcriptional compensation when *apiAT5-3* is deleted.

**Fig 3.**
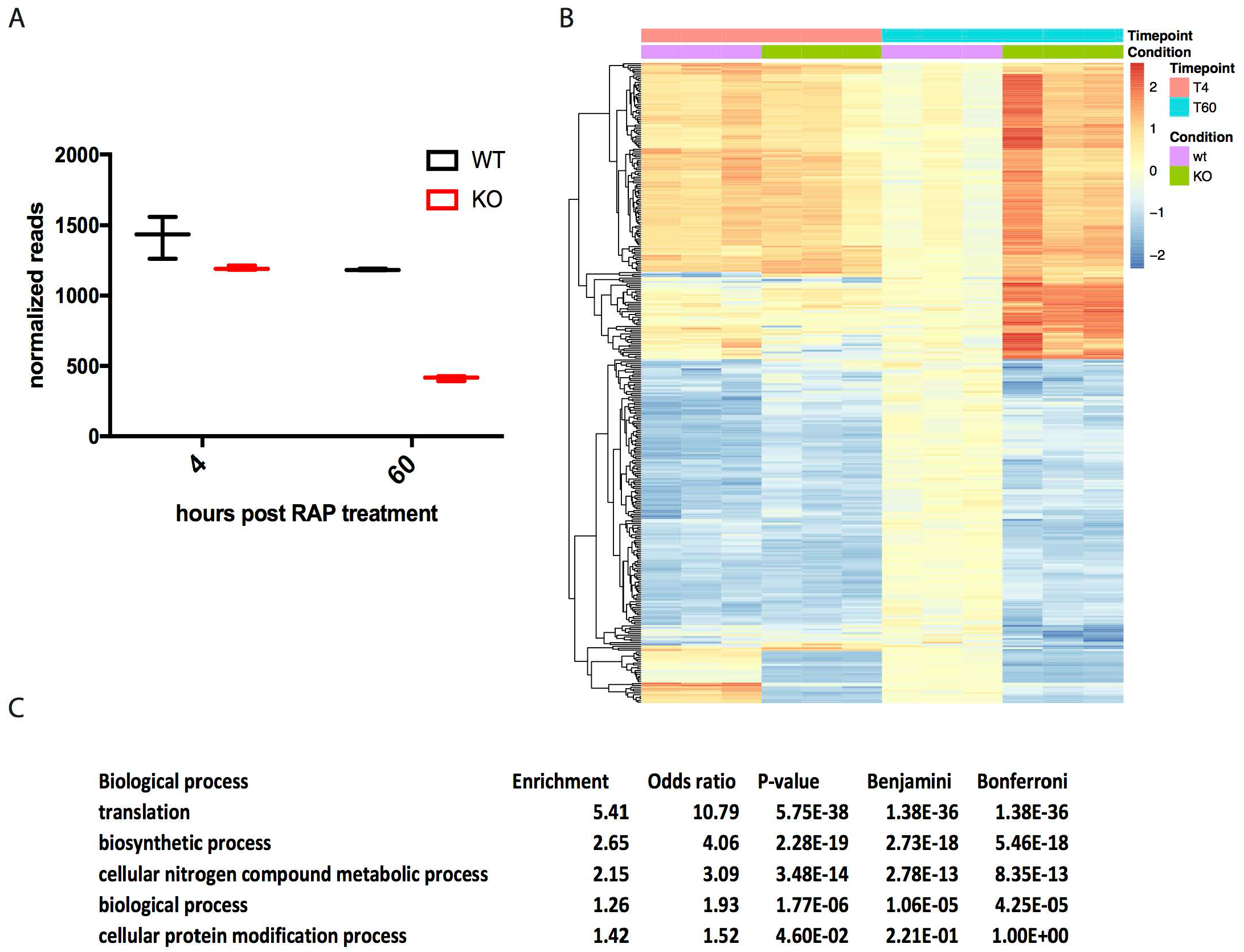
Δ*apiAT5-3* parasites display a transcriptional response related to amino acid starvation. **(A)** Extracted reads for recodonised *apiAT5-3* from RNA sequence data show a significant reduction of *apiAT5-3* transcripts in RAP-treated ApiAT5-3_loxP lines 60 hrs post RAP-treatment compared to RAP-treated ApiAT5-3-loxP^dDiCre^ parasites. **(B)** Heatmap of genes that change significantly (adjusted p <0.05) in transcript read number between WT and Δ*apiAT5-3* 60 hrs post addition of RAP. **(C)** Gene ontology term enrichment shows that genes involved in translation processes are significantly enriched among the differentially expressed genes 60 hrs post RAP-treatment.

Collectively these data show that ApiAT5-3 is an essential protein that is required for intracellular replication. Its depletion leads to a complete arrest in growth which is not accompanied by a substantial stress response, but rather modest signs of translational stress.

### Mutation of S56 to alanine, but not a phosphomimetic leads to a reduction in fitness

Having established that ApiAT5-3 is essential for the lytic cycle, we next sought to examine the role of *Tg*CDPK3-mediated phosphorylation in ApiAT5-3 function. To do this, we complemented ApiAT5-3_loxP parasites with either WT ApiAT5-3, or variants where S56 is mutated to alanine (S56A) or to aspartic acid (S56D). To prevent possible differences in growth between the parasite lines due to differential expression of the complementation constructs, we inserted each into the *uprt* locus by double homologous recombination, under control of the endogenous promoter (Fig. 4A). Complementation into the *uprt* locus was verified by PCR (Fig. 4B). The complementation constructs also carried a C-terminal HA epitope tag to verify correct trafficking to the plasma membrane. Immunofluorescence displayed correct trafficking in all variants (Fig. 4C).

**Fig 4.**
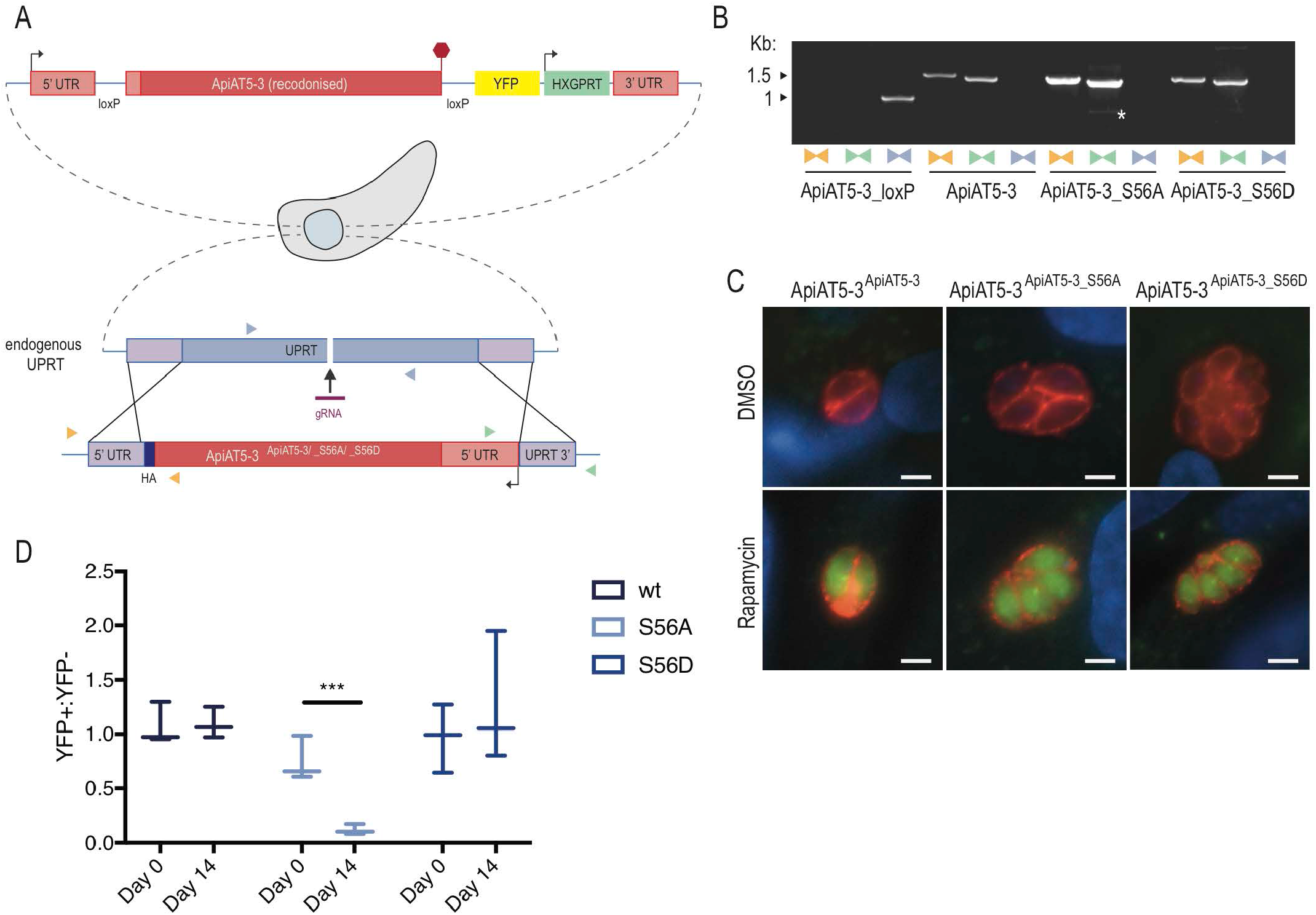
Δ*apiAT5-3*^ApiAT5-3_S56A^ demonstrates a fitness defect. **(A)** Generation of the ApiAT5-3^ApiAT5-3/_S56A/_S56D^ complementation lines. PAM indicated by black arrow. Primer pairs represented by coloured triangles. **(B)** PCR analysis shows correct integration of the ApiAT5-3-loxP construct at both the 3’ and 5’ ends and a loss of *uprt*. White * = non-specific band **(C)** IFA of ApiAT5-3^ApiAT5-3/_S56A/_S56D_HA^ expressing parasites shows that ApiAT5-3 is correctly trafficked to the periphery of the intracellular tachyzoite in both the presence (DMSO) and absence (Rapamycin) of the endogenous *apiAT5-3*. Red = HA. Green = YFP, indicating correct excision of the endogenous *apiAT5-3*. Scale bar 10 μm. **(D)** Growth competition assay by flow cytometry shows that Δ*apiAT5-3*^ApiAT5-3_S56A^ parasite growth is reduced relative to the non-excised ApiAT5-3^ApiAT5-3_S56A^ line. Statistical analysis using multiple comparison, 2-way ANOVA of mean ratio to day 0 normalised to 1. ***p <0.001, n = 3.

To compare fitness between the WT, the phosphomimetic (S56D), and the phosphomutant (S56A) complemented lines in the absence of *apiAT5-3*, we deleted the endogenous copy using RAP-treatment. This results in parasite strains that solely rely on the complemented copy of the gene. We confirmed correct excision of *apiAT5-3* by virtue of YFP expression post RAP-treatment, and PCR analysis (Fig. 4C, Fig. S2A). RAP-treated parasite lines were viable and allowed us to isolate clones by limiting dilution, all of which restored growth in plaque assays (Fig. S2B). This shows that i) complementation of *apiAT5-3* by expression at the *uprt* locus fully restores ApiAT5-3 function and ii) that neither the introduction of phosphomimetics nor phosphomutants of S56 are lethal to parasite growth. This is not surprising as deletion of *Tg*CDPK3, the kinase putatively responsible for ApiAT5-3 phosphorylation during egress, does not lead to a severe growth phenotype. Accordingly, phosphomutants would not be expected to display drastic differences in growth. We therefore performed competition assays in which we compared growth of YFP expressing complementation lines that fully rely on the complementation variant for growth (Δ*apiAT5-3*^ApiAT5-3/S56A/S56D^) mixed in a 1:1 ratio with their non-excised, colourless counterpart (ApiAT5-3^ApiAT5-3/S56A/S56D^). Using the ratio of 4’,6-diamidino-2-phenylindole (DAPI) stained parasites (DAPI labels the DNA of all parasites) and YFP expressing parasites (YFP is expressed only in the complementation lines) we followed growth over 14 days in biological triplicates. While Δ*apiAT5-3*^ApiAT5-3^ parasites showed no difference in growth compared to their WT control, Δ*apiAT5-3*^ApiAT5-3_S56A^ was reduced by 84.0% after 14 days (Fig. 4D). Strikingly, Δ*apiAT5-3*^ApiAT5-3_S56D^ was not outcompeted and grew at similar levels to the WT control. Collectively these data indicate that phosphorylation of S56 while not essential, is important for intracellular growth.

### ApiAT5-3 is a primary transporter of tyrosine, but not branched chain amino acids

The predicted homology of ApiAT5-3 to a BCAA transporter and the profound up-regulation of the BCKDH complex in Δ*cdpk3* parasites suggested a direct role for ApiAT5-3 in BCAA transport. To test this, we expressed *apiAT5-3* in the heterologous expression system, *X. laevis* oocytes. Concurrently with our study, data were presented that ApiAT5-3 may be a tyrosine transporter (Giel van Dooren, personal communication and pre-published in BioRx (19)). We therefore tested BCAA import and replicated the tyrosine uptake capacity of ApiAT5-3 in oocytes expressing WT ApiAT5-3 (Fig. 5A). Measuring unidirectional influx, we observed a significant (4.0-fold) increase in the uptake of ^14^C-tyrosine into ApiAT5-3-expressing oocytes compared to either water-injected or uninjected control oocytes under the conditions tested, consistent with results from (19). We also observed moderate ApiAT5-3-dependent phenylalanine influx, but not for the BCAA valine (Fig. S3A), suggesting that, while ApiAT5-3 is capable of tyrosine transport, it is unlikely to be a major BCAA transporter.

**Fig 5.**
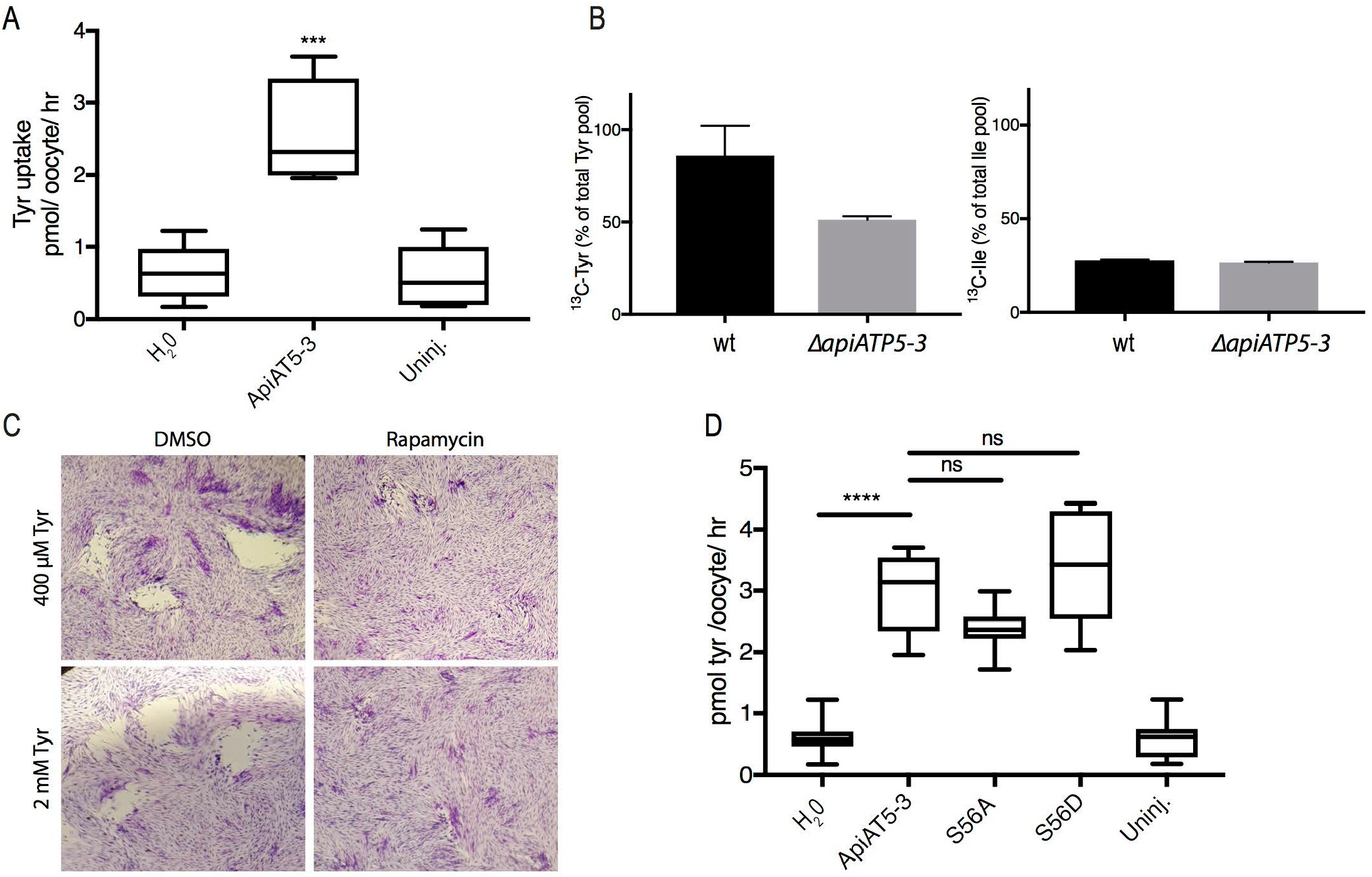
Functional analysis of the ApiAT5-3 transporter. **(A)** *X. laevis* oocytes expressing ApiAT5-3 demonstrate a significant increase in ^14^C-L-tyrosine uptake. 10 oocytes per experiment. Analysis carried out using a two-tailed, paired, Student’s t-test. ***p <0.001 Box plots show mean, 1^st^ and 3^rd^ quartile and SD, n= 5. **(B)** Extracellular, RAP-treated, ApiAT5-3_loxP tachyzoites, labelled with ^13^C-L-tyrosine or ^13^C-L-isoleucine, display a marked decrease in tyrosine but not isoleucine import, relative to WT. n=2. **(C)** Plaque assay shows no rescue of growth of RAP-treated ApiAT5-3_loxP on addition of excess (2 mM) L-tyrosine. **(D)** *X. laevis* oocytes expressing *apiAT5-3_S56A* demonstrate a modest but insignificant reduction in ^14^C-L-tyrosine uptake relative to *apiAT5-3* and *apiAT5-3_S56D*. 10 oocytes per experiment ****p <0.0001, ns = non-significant. Analysis carried out using multiple comparison, one-way ANOVA. Box plots show mean, 1^st^ and 3^rd^ quartile and SD, n= 5.

To verify the role of ApiAT5-3 in tyrosine transport in our conditional KO parasites, we measured intracellular ^13^C-tyrosine levels in RAP-treated Δ*apiAT5-3*^ApiAT5-3^ (WT) and ApiAT5-3_loxP (KO) parasites (74 hrs post excision), after 1 hr in the presence of growth media containing ^13^C-tyrosine. In an analogous manner, we also measured ^13^C-isoleucine uptake in order to verify if ApiAT5-3 is also a BCAA transporter. Δ*apiAT5-3*^ApiAT5-3^ was used instead of DMSO-treated ApiAT5-3_loxP to control for any potential effects of rapamycin on parasite metabolism. This analysis verified a reduction of ^13^C-labelled tyrosine uptake (40.5% compared to Δ*apiAT5-3*^ApiAT5-3^), but not isoleucine uptake (4.3% compared to Δ*apiAT5-3*^ApiAT5-3^) (Fig. 5B). We also measured the intracellular abundance of all detectable amino acids when labelling with ^13^C-tyrosine. We observed a reduction of intracellular tyrosine abundance (63.2%) in the Δ*apiAT5-3*^ApiAT5-3^ cells (as expected), but not phenylalanine which was slightly increased in relative abundance (17.45%), suggesting that while ApiAT5-3 is able to transport phenylalanine in oocytes, it is not the major phenylalanine transporter in *T. gondii* (Fig. S3B). It is important to note that our metabolome analysis was performed at the end of cycle 2 after RAP-treatment, when Δ*apiAT5-3*^ApiAT5-3^ parasites are still viable but start to display a reduction of growth (Fig. 2e). Therefore, we predict that low levels of ApiAT5-3^ApiAT5-3^ present at this stage are responsible for the residual transport of tyrosine. Interestingly, we also observed a reduction in intracellular aspartate (38.5%) and glycine (28.3%) in Δ*apiAT5-3*^ApiAT5-3^ cells (Fig. S3B). Since *T. gondii* is not known to be auxotrophic for these amino acids we reasoned that the observed death phenotype is unlikely caused by a defect in glycine or aspartate import, and instead focussed our subsequent analysis on tyrosine. We also observed an increase in the abundance of glutamine, valine, isoleucine and proline, indicating potential wider metabolic effects.

To test whether exogenous tyrosine can complement the loss of ApiAT5-3 we grew parasites in media with 5× the normal amount of tyrosine (2 mM). Despite various attempts to restore normal growth or generate Δ*apiAT5-3* clonal lines, we could not obtain viable parasites in high tyrosine (Fig. 5C). In other organisms, phenylalanine can be converted into tyrosine. Therefore, we tested whether phenylalanine supplementation (2 mM) can rescue the growth phenotype of *apiAT5-3* KO parasites. No growth rescue could be observed (data not shown). Together, these results suggest that ApiAT5-3 is the only transporter of tyrosine in *T. gondii* and that phenylalanine cannot be readily converted into tyrosine in *T. gondii* parasites.

To identify whether differences in phosphorylation state of ApiAT5-3 affected tyrosine transport, we performed isotopic tyrosine labelling of extracellular parasites using the Δ*apiAT5-3*^ApiAT5-3_S56A^ and Δ*apiAT5-3*^ApiAT5-3_S56D^ phosphomutant strains. No significant differences could be observed in these assays indicating that, if tyrosine import is affected, differences may be obstructed by the intrinsic variability of the assay (data not shown).

The fitness phenotype of mutating S56 to alanine is modest in a single lytic cycle and thus not predicted to have substantial impact on the import of tyrosine. Because we could not measure differences in tyrosine import in our phosphorylation site mutants we turned to heterologous assays where we expressed ApiAT5-3 WT, S56A and S56D in *X. laevis* oocytes and tested tyrosine uptake. Although there was a trend towards a reduction in tyrosine uptake in the S56A expressing oocytes (average 19.5% reduction in S56A relative to WT ApiAT5-3 expressing oocytes), this difference was not statistically significant (Fig. 5D). The S56D expressing oocytes display a marginal increase in tyrosine uptake of 14.0%, that again was not statistically significant.

Collectively these data show that conditional deletion of ApiAT5-3 causes a lethal reduction in tyrosine import that cannot be compensated for by amino acid supplementation. We also show that while phosphorylation of ApiAT5-3 at S56 is not required for tyrosine transport, mutating that site to alanine does lead to a subtle decrease in tyrosine transport in a heterologous assay.

## Discussion

*Tg*CDPK3 has previously been implicated in controlling distinct biological processes such as gliding motility and metabolism. How these are linked, however, has been unclear. Mutants that display only IIE and IID phenotypes have been identified (9,26), arguing that *Tg*CDPK3 may be an upstream regulator of both processes. Here we show that upon activation by the Ca^2+^ ionophore A23187, *Tg*CDPK3 leads to an increase in ApiAT5-3 phosphorylation at S56. This occurs at the same time, and to a similar intensity, as other previously identified targets of *Tg*CDPK3 (e.g. serine 21/22 of MyoA) and other kinases involved in signalling (e.g. *Tg*CDPK1). MyoA and ApiAT5-3 are both located at, or close to, the plasma membrane. It is conceivable that, upon activation, *Tg*CDPK3 phosphorylates proteins at the plasma membrane, some of which are important for motility and others (e.g. transporters) that prepare the parasite for the extracellular milieu. In this study we have identified a function for the phosphorylation of S56, which becomes rapidly phosphorylated prior to egress in a *Tg*CDPK3-dependent manner. Interestingly, ApiAT5-3 possesses several other phosphorylation sites in its N-terminus, aside from S56, that either do not change in phosphorylation state or, in the case of S14, appear dephosphorylated during induced Ca^2+^ signalling. How S14 dephosphorylation and S56 phosphorylation are controlling ApiAT5-3 function requires further investigation, however it is evident that mutating S56 to a non-phosphorylatable residue substantially reduces parasite fitness. Phosphorylation of transporters has been shown to regulate affinity, specificity and flux of cargo (27–32). Accordingly, the observed fitness cost in S56A mutants could be indicative of a reduction in tyrosine import, for which the parasite is auxotrophic (33).

While we observed a clear role for ApiAT5-3 in tyrosine transport, expression of the ApiAT5_S56A phosphomutant only led to a modest reduction in tyrosine import in oocytes, which was not statistically significant. One explanation could be that the effect on tyrosine import is beyond the limit of detection in our assays. We observed an 84.0% reduction in growth of Δ*apiAT5-3*^ApiAT5-3_S56A^ mutants compared to WT parasites over 14 days. This translates into a reduced replication rate of ~6.74% per 24 hrs. Tyrosine uptake assays in oocytes, as well as in extracellular parasites are, for reasons of cell viability, performed in <1 hr, and may therefore not pick up the subtle differences predicted to occur if tyrosine is the only limiting factor in these assays. Furthermore, as indicated above, ApiAT5-3 shows a complex phosphorylation pattern at its N-terminus and other phosphorylation sites could contribute to ApiAT5-3 regulation. Mimicking these conditions in a heterologous assay, or in parasites, without prior knowledge of the abundance of the phosphorylation on each of these residues becomes hard to interpret and should be the subject of further studies.

Apart from regulating amino acid transport, phosphorylation of transporters has also been shown to regulate trafficking to the surface (34–36). However, a role for S56 in trafficking is less likely for two reasons: i) we did not observe any obvious defects in surface translocation of the transporter in parasites and ii) *Tg*CDPK3 phosphorylates S56 shortly before, or during egress, at which state the transporter is already on the surface. If S56 phosphorylation was important for surface translocation, we would expect this to occur at an earlier stage. However, we cannot exclude the possibility that minor differences in trafficking capacity impact tyrosine transport, resulting in the growth phenotype.

Whatever the molecular explanation for the phenotype, it is evident that *Tg*CDPK3-mediated phosphorylation of ApiAT5-3 is important for parasite fitness.

Interestingly we haven’t been able to rescue the effect of *apiAT5-3* loss in our conditional KO parasites through growth in high tyrosine concentrations. These results differ from those in Parker, K., *et al* whereby growth can, at least partially, be rescued in high tyrosine medium (19). One mechanism to counter low tyrosine levels that most organisms possess, is the ability to convert phenylalanine into tyrosine, via the enzyme phenylalanine-4-hydroxylase (AAH). However, the lack of tyrosine import in ApiAT5-3 conditional KO’s cannot be overcome in the presence of high levels of phenylalanine, indicating that this pathway is not available. Indeed, both isoforms of AAH (AAH1 and AAH2) have recently been shown to be secreted into the host cell during *Toxoplasma* infection, where they would be unable to rescue a tyrosine transporter defect in the plasma membrane (33,37). Another explanation as to why Parker, K., *et al*. have successfully rescued the effects of *apiAP5-3* deletion through addition of excess tyrosine, could be via the upregulation of alternative transporters. Although a small amount of tyrosine appears to be imported in our *apiAT5-3* KO line (Fig. 5B), this is likely due to the presence of residual ApiAT5-3 protein in the plasma membrane after RAP-treatment. Along with our inability to rescue growth upon tyrosine supplementation, we conclude that this residual tyrosine import is unlikely due to an alternative transporter. Further to this, our transcriptomic analysis argues against a rapid sensing and transcriptional compensation for the lack of tyrosine import, so if upregulation of alternative transporters occurs, it will be a slow process. Another explanation may be that slight differences in the genetic background or passage history, and potential epigenetic changes in the parental strains, leads to a difference in capacity for amino acid transport. There is some indication that this may be the case as, in our metabolomics experiments, the ApiAT5-3 deletion showed reduced levels of glycine and aspartic acid in addition to tyrosine, while in Parker et al., other amino acids where observed to be less abundant in addition to tyrosine. We also saw an increase in abundance of some amino acids that differ from Parker, K *et al*. It may be interesting in the future, to compare our *apiAT5-3* KO with that of Parker, K., *et al*. and identify any compensatory mechanisms the parasites can use to adjust to tyrosine starvation. Interestingly, deletion of *apiAT5-3* leads to a growth arrest that is not accompanied by major transcriptional responses. This is reminiscent of the hibernation state in *Plasmodium falciparum*, whereby depletion of isoleucine, an essential amino acid for this parasite, leads to arrest in growth without a major stress response (38). This would suggest that translational arrest may be a common response among apicomplexan parasites during amino acid starvation.

In summary we show that ApiAT5-3, a novel *T. gondii* tyrosine transporter, is rapidly phosphorylated in a *Tg*CDPK3 dependent manner at S56 prior to, and during egress from the host cell. The phosphorylation of S56 appears important for parasite fitness. These results, together with previous studies, support the notion that *Tg*CDPK3 simultaneously targets several proteins in, or at, the plasma membrane, controlling very divergent biological processes, such as motility and metabolism. The phenotypes observed in Δ*cdpk3* parasites may therefore be an accumulation of effects on various proteins, which is likely true for other kinases as well.

## Materials and Methods

### Parasite culture

*T. gondii* parasites were cultured in a confluent monolayer of human foreskin fibroblasts (HFFs) maintained in Dulbecco’s Modified Eagle Medium (DMEM), GlutaMAX supplemented with 10% foetal bovine serum, at 37°C and 5% CO_2_.

### Plasmid and parasite strain generation

A comprehensive list of primers and parasite lines used throughout this study are described in S1 and S2 Tables respectively. To generate the epitope tagged ApiAT5-3::HA line, the *apiAT5-3* gene and associated 5’ UTR were PCR-amplified from RH *T. gondii* gDNA using the primers 1 and 2 and cloned using Gibson assembly (39) into pGRA::HA::HPT (40), linearised with HindIII and NcoI. 25 μg of the pGRA::ApiAT5-3::HA vector was transfected into RH Δ*hxgprt* parasites as previously described (41). 16-20 hrs after transfection, transgenic parasites were selected using 25 μM mycophenolic acid (MPA) and 50 μg/ml xanthine (XAN). To generate the ApiAT5-3_loxP conditional KO lines, the *apiAT5-3* 5’UTR was first PCR-amplified from gDNA with primers 3 and 4. This PCR product was inserted, along with the synthetic DNA constructs *loxP_apiAT5-3_loxP_yfp* and *loxP*(−100)_*apiAT5-3* (see S1 Table for full sequences), by Gibson assembly into pG140::Actin::YFP (42) that had been PCR-amplified using primers 5 and 6 to remove the actin gene. 2 μg of the subsequent pG140::ApiAT5-3_loxP::YFP plasmid was linearised with ScaI and co-transfected into RH *Δku80Δhxgprt* with pSag1::Cas9-U6::dbl-sgApiAT5-3, in a molar ratio of 1:10. The pSag1::Cas9-U6::dbl-sgApiAT5-3 vector was generated by PCR-amplification of the pSag1::Cas9-U6 (43) vector using primers 7 and 8 to insert the 5’ gRNA (gRNA 1) and 9 and 8 to insert the 3’ gRNA (gRNA 2), prior to re-ligation with T4 DNA Ligase (New England Biolabs M0202). gRNA 1 was then amplified using primers 10 and 11 and Gibson cloned into the pSag1::Cas9-U6::sg2ApiAT5-3 that had been linearised with KpnI and XhoI as per (25). Transgenic parasites were selected using MPA/XAN as described for pGRA::ApiAT5-3::HA. 5’ and 3’ integration was confirmed using primer pairs 12 and 13, and 14 and 15 respectively. Absence of WT *apiAT5-3* was confirmed using primers 16 and 17. DiCre-mediated *apiAT5-3* excision was induced with the addition of 50 nM RAP to ApiAT5-3_loxP parasites for 4 hrs. Excision was confirmed using primers 13 and 16. To introduce an ectopic copy of *apiAT5-3* into the *uprt* gene locus, the *apiAT5-3* gene, and associated 5’ UTR, were PCR-amplified from gDNA using primers 18 and 19 which was then inserted into the BamHI/ PacI digested pUPRT::DHFR-D vector (Addgene plasmid #58258 (43)) using Gibson assembly. To generate the pUPRT::ApiAT5-3_S56A::HA and pUPRT::ApiAT5-3_S56D::HA vectors, the pUPRT::ApiAT5-3::HA vector was modified by site directed mutagenesis using the primers 20 and 21 (S56A) or 22 (S56D). pUPRT::ApiAT5-3::HA, pUPRT::ApiAT5-3_S56A::HA and pUPRT::ApiAT5-3_S56D::HA were linearised with PciI prior to the co-transfection of 2 μg into RH *Δku80Δhxgprt* ApiAT5-3_loxP with pSag1::Cas9-U6::sgUPRT (Addgene 54467 (43)) in a molar ratio of 1:10. Transgenic parasites were selected by the addition of 5 μM 5’-fluo-2’-deoxyuridine to culture medium, 16-20 hrs post-transfection. Integration into the genome was confirmed using primer pairs 24 and 25, and 26 and 27 respectively. Absence of *uprt* was confirmed using primers 28 and 29.

### Western blotting and immunofluorescent imaging

For Western blot analysis, intracellular parasites were lysed in Laemmli buffer (60 mM Tris-HCl pH6.8, 1% SDS, 5% glycerol, 5% b-mercaptoethanol, 0.01% bromophenol blue) and heated to 37°C for 30 mins prior to separation on a 10% sodium dodecyl-polyacrylamide gel. Proteins were transferred onto a nitrocellulose membrane prior to blocking in 3% milk, 0.1% Tween-20 PBS. HA-tagged ApiAT5-3 was detected using rat anti-HA (1:500), followed by goat anti-rat horseradish peroxidase-conjugated secondary antibody (1:2500).

IFA’s were performed on intracellular parasites grown in HFFs on glass coverslips. 1×10^5^ parasites were seeded 24 hrs prior to fixation with 3% formaldehyde (FA). PBS 0.1% Triton X-100 was added to the fixed cells for 10 mins prior to blocking with 3% bovine serum albumin in PBS for 1 hr. ApiAT5-3::HA was visualised using rat anti-HA (1:500) followed by addition of Alexa488 conjugated donkey anti-rat secondary antibody (1:2000) and DAPI, 5 μg/ml.

### Plaque assay and amino acid complementation

For plaque assay analysis, 150 parasites were seeded on confluent HFF monolayers, grown in 24-well plates, and left undisturbed for 5 days, before fixing with chilled methanol and staining with 0.1% crystal violet. To assess growth in excess tyrosine, plaque assays were repeated either at normal tyrosine levels (400 μM L-tyrosine disodium salt; as per Gibco manufacturer) or in DMEM supplemented with 2mM L-tyrosine disodium salt (dissolved for 1 hour at 50 °C). To ensure tyrosine had successfully dissolved samples of the media were analysed by GC-MS as previously described (18).

### Replication assay

2×10^4^ ApiAT5-3_loxP parasites were seeded in triplicate on confluent HFFs in both culture flasks and glass bottom 8-well imaging plates and left to invade for 1 hour prior to treatment with 50 nM RAP or equivalent volume DMSO, for 4 hrs. Parasites were imaged at 24, 36 and/or 48 hrs and split at 36 hrs into new flasks and imaging wells for the subsequent replication cycle. At each time point parasites were fixed in 3% FA and imaged on a Nikon Eclipse Ti-U inverted fluorescent microscope. Parasites/vacuole were counted manually from 5 fields of view at 20× magnification using the Nikon NIS-Elements software.

### Live cell microscopy

ApiAT5-3_loxP parasites were treated with RAP or DMSO as previously described. 36 hrs into cycle 2 post RAP-treatment parasites were syringe lysed and seeded in glass bottom, 8-well imaging plates in a 1:1 ratio with RH Tom parasites. After a further 29 hrs, live parasites were imaged on a Nikon Eclipse Ti-U inverted fluorescent microscope every 30 mins for the next 30 hrs, in a temperature-controlled chamber at 37 °C and 5% CO_2_. Images were analysed using the Nikon NIS-Elements software.

### Ionophore induced egress and death assays

ApiAT5-3_loxP parasites were seeded in 96-well imaging plates at a MOI of 0.5, 36 hrs post RAP/DMSO-treatment. Ionophore induced egress assays were performed in triplicate at 37 °C in Ringers buffer (155 mM NaCl, 3 mM KCl, 2 mM CaCl_2_, 1 mM MgCl_2_, 3 mM NaH_2_PO_4_, 10 mM HEPES, 10 mM glucose) 30 hrs later. The parasites were incubated with 8 μM Ca^2+^ ionophore A23187 for 0, 0.5, 1, 1.5, 2, 2.5, 3, 3.5, 4 and 5 mins prior to the addition of 16% FA to a final concentration of 3% for 15 mins. Wells were subsequently washed with PBS and stained with 5 μg/ml DAPI. Automated image acquisition of 25 fields per well was performed on a Cellomics Array Scan VTI HCS reader (Thermo Scientific) using a 20× objective. Image analysis was performed using the Compartmental Analysis BioApplication on HCS Studio (Thermo Scientific).

### Competition assays and flow cytometry

5×10^6^ ApiAT5-3^ApiAT5-3^, ApiAT5-3^ApiAT5-3_S56A^ and ApiAT5-3^ApiAT5-3_S56D^ parasites were mixed in a 1:1 ratio with Δ*apiAT5-3*^ApiAT5-3^, Δ*apiAT5-3*^ApiAT5-3_S56A^ and Δ*apiAT5-3*^ApiAT5-3_S56D^ respectively. 5×10^4^ parasites were added to fresh HFF monolayers before spinning the rest of the sample at 72 × g to remove host cell debris for 1 min. The supernatant was spun at 2049 × g for 5 mins. The pellet was fixed for 10 mins in 3% FA, washed in PBS and stained with 5 μg/ml DAPI. The sample was washed and resuspended in PBS before running on a flow cytometer. All parasites were gated on DAPI fluorescence to prevent results being skewed by remaining unstained host cell debris. The proportion of DAPI (+); YFP (+) (representing Δ*apiAT5-3*^ApiAT5-3/S56A/S56D^) compared to DAPI (+); YFP (−) (representing ApiAT5-3^ApiAT5-3/S56A/S56D^) was calculated. The process was repeated 14 days later for comparison to day 0.

### Oocyte maintenance and radiotracer uptake assays

ApiAT5-3, ApiAT5-3_S56A and ApiAT5-3_S56D were PCR amplified from Δ*apiAT5-3*^ApiAT5-3^, Δ*apiAT5-3*^ApiAT5-3_S56A^ and Δ*apiAT5-3*^ApiAT5-3_S56D^ cDNA, respectively, using primers 30 and 31 to add a region of homology to the XkbN plasmid at the 5’ end and a HA tag to the 3’ end of each gene. These fragments were then amplified with primers 32 and 33 to add a 3’ XkbN homology overhang. These resulting fragments were inserted by Gibson assembly into the XkbN1_*Pf*HT (a version of pSPGT1 (44) with a NotI site added to the MCS, provided by Ksenija Slavic) which had been digested with BglII and NotI, to remove the *Pf*HT gene. The resulting XkbN_ApiAT5-3, XkbN_ApiAT5-3_S56A and XkbN_ApiAT5-3 _S56D plasmids were linearised with XbaI for in vitro transcription using the Thermo Fisher mMessage mMachine transcription kit. Stage V to VI defolliculated *X. laevis* oocytes were obtained commercially (Ecocyte Biosciences) for subsequent functional transport studies. Oocytes were microinjected with cRNA (20 to 40 ng in 30 nl of water) encoding *apiAT5-3* template or with a comparable amount of diethylpyrocarbonate-treated water. The oocytes were maintained at 18 °C in oocyte Ringer 2 buffer (82.5 mM NaCl, 2.5 mM KCl, 1.5 mM CaCl_2_, 1mM Na_2_HPO_4_, 1 mM MgCl_2_ and 5 mM HEPES) and used for transport studies 72 hrs after cRNA injection. Transport measurements were performed at room temperature on groups of 10 oocytes in ND96 medium (96 mM NaCl, 2 mM KCl, 2 mM CaCl_2_, 1 mM MgCl_2_ and 5 mM HEPES) containing 1 μM radiolabelled U-^14^C-L-tyrosine (specific activity of 486 mCi/mmol; Perkin Elmer), U-^14^C-L-phenylalanine (specific activity of 508 mCi/mmol; Perkin Elmer) or U-^14^C-L-valine (specific activity of 271 mCi/mmol; Perkin Elmer). Transport was measured at 10 min, over which time uptake of L-tyrosine is linear (19). Each result was confirmed by at least 3 independent experiments.

### Metabolite labelling and extraction

ApiAT5-3-loxP, Δ*apiAT5-3*^ApiAT5-3^, Δ*apiAT5-3*^ApiAT5-3_S56A^ and Δ*apiAT5-3*^ApiAT5-3_S56D^ parasites were treated in triplicate with 50 nM RAP and, at the end of the first cycle, seeded in 15 cm culture flasks. Stable isotope labelling (1 hr) of extracellular parasites with 0.8 mM U-^13^C-L-tyrosine or 4 mM U-^13^C-L-isoleucine, metabolite extraction and subsequent GC-MS analysis were all performed as per (18), on an Agilent GC-MSD (7890B-5977A). Data analysis was carried out using GAVIN software (45).

### RNA sequencing analysis

*T. gondii* RNA was extracted as per the Qiagen RNA-easy mini kit user handbook (#74104) from ~5×10^6^ ApiAT5-3_loxP or ApiAT5-3_loxP^dDiCre^ parasites at 0, 4 and 60 hrs post RAP-treatment. Analysis was performed in triplicate. The FASTQ files were aligned using Bowtie 2 (46) to Ensembl Protist’s release 35 of *T. gondii* (ToxoDB-7.1). They were then quantified using RSEM before being processed using Bioconducor (47). We used DESeq2 (48) to account for gene length and library size, and to test for the interaction between treatment and time point to generate the differential genelist. We corrected for multiple testing using the Benjamini-Hochberg procedure for false discovery rates. To validate the recodonised transcript, we both re-aligned to a custom genome rebuilt to include the novel sequence, and also used a pseudo-alignment approach to quantify purely the reads associated with the novel sequence (49).

## Acknowledgements

We thank all members of the Treeck laboratory for critical discussions. We thank Giel Van Dooren and Sebastian Lourido for sharing unpublished data. We thank members of the following Science Technology platforms at the Francis Crick Institute: Bioinformatics, Advanced Sequencing, Peptide Synthesis, Proteomics and Flow Cytometry.

## Supporting information

**S1 Fig. ApiAT5-3_loxP parasites that survive RAP-treatment retain the *apiAT5-3* gene**.

PCR analysis using primers spanning the floxed *apiAT5-3* gene show that the small proportion of non-excised parasites present after RAP-treatment outgrow the excised Δ*apiAT5-3* parasites within 2 weeks.

**S2 Fig. Verification of ApiAP5-3 complementation lines**.

**(A)** Addition of RAP to the complemented parasite lines leads to correct recombination of the loxP sites and deletion of the endogenous *apiAT5-3* gene. **(B)** Plaque assay showing restoration of plaquing efficiency upon RAP-treatment of the ApiAT5-3-loxP line complemented with the WT or phosphomutant versions of the gene.

**S3 Fig. Analysis of ApiAP5-3 transport function**.

**(A)** *X. laevis* oocytes expressing *apiAT5-3* demonstrate an increase in ^14^C-L-phenylalanine uptake but no significant ^14^C-L-valine uptake. 10 oocytes per experiment. Analysis carried out using a two-tailed, paired, Student’s t-test. *p <0.05, ns = non-significant. n= 2. **(B)** Relative abundance of amino acids in RAP-treated ApiAT5-3-loxP (KO) relative to Δ*apiATP5-3*^ApiAT5-3^ (WT), shows that tyrosine is the most significantly reduced upon loss of ApiAT5-3. **(C)** Tyrosine abundance in normal DMEM compared to DMEM supplemented with 2 mM tyrosine.

**S1 Movie. Live video microscopy of Δ*apiAT5-3* parasites**

Live video microscopy of ApiAT5-3_loxP parasites 29 hrs into the 3^rd^ lytic cycle post DMSO- **(A)** or RAP- **(B)** treatment. Red = WT RH Tom, colourless parasites = non-excised ApiAT5-3_loxP, green parasites = YFP expressing *apiAT5-3* KO.

**S1 Table. Primers and synthetic DNA sequences used throughout this study.**

**S2 Table. *Toxoplasma gondii* strains generated throughout this study.**

**S3 Table. RNA sequencing analysis of *apiAT5-3* conditional KO**

A list of all genes displaying log^2^ fold change in the RAP-treated Δ*apiAT5-3*^ApiAT5-3^ compared to ApiAT5-3_loxP parasites, 4 and 60 hrs after RAP-treatment.

